# Sensitivity of Olfactory Sensory Neurons to food cues is tuned to nutritional states by Neuropeptide Y signalling

**DOI:** 10.1101/573170

**Authors:** Tarun Kaniganti, Ajinkya Deogade, Aditi Maduskar, Arghya Mukherjee, Akash Guru, Nishikant Subhedar, Aurnab Ghose

**Affiliations:** Indian Institute of Science Education and Research (IISER) Pune, Dr Homi Bhaba Road, Pune 411008, INDIA

**Keywords:** Neuropeptide Y, Olfactory sensitivity, Olfactory Sensory Neurons, nutritional state, neuromodulation

## Abstract

**Background:** Modulation of sensory perception by homeostatic feedback from physiological states is central to innate purposive behaviours. Olfaction is an important predictive modality for feeding-related behaviours and its modulation has been associated with hunger-satiety states. However, the mechanisms mapping internal states to chemosensory processing in order to modify behaviour are poorly understood.

**Results:** In the zebrafish olfactory epithelium, a subset of olfactory sensory neurons (OSNs) and the terminal nerve projections express neuropeptide Y (NPY). We find that NPY signalling in the peripheral olfactory system of zebrafish is correlated with its nutritional state and is both necessary and sufficient for the olfactory perception of food related odorants. NPY activity dynamically modulates the microvillar OSN activation thresholds and acts cooperatively with amino acid signalling resulting in a switch-like increase in OSN sensitivity in starved animals. We suggest that cooperative activation of phospholipase C by convergent signalling from NPY and amino acid receptors is central to this heightened sensitivity.

**Conclusions:** This study provides ethologically relevant, physiological evidence for NPY signalling in peripheral modulation of OSN sensitivity to food-associated amino acid cues. We demonstrate sensory gating directly at the level of OSNs and identify a novel mechanistic framework for tuning olfactory sensitivity to prevailing energy states.

## INTRODUCTION

Modulation of sensory perception by homeostatic feedback is critical for survival. Gating of sensory inputs by internal states, such as energy availability, enables the animal to issue measured responses, like food-seeking and consumptive behaviours. Olfaction is an important sensory modality in food seeking behaviour providing predictive information on the availability, quality and associated reward salience. Nutritional state-dependent modulation of olfactory perception has been reported in a number of species, though the underlying mechanisms are not well understood (Palouzier-Paulignan *et al.*, 2012; Sengupta, 2013). Additionally, dysfunctional olfactory responses are reported in obese animals, including humans, highlighting the importance of this sensory modality in health and disease (Aime *et al.*, 2007; Stafford *et al.*, 2013).

Activity patterns of neuromodulators generate molecular representations of internal states and modify neuronal properties and circuit dynamics to effect homeostatic adjustments. Neuromodulatory inputs may impinge on multiple levels of sensory processing, including the peripheral sensory circuitry. In the olfactory system, ghrelin, orexins, insulin, leptin, GnRH, FMRF-amide, cholecysteokinin and endocannabinoids are known to influence olfactory sensitivity (Eisthen *et al.*, 2000; Park & Eisthen, 2003; Hardy *et al.*, 2005; Baly *et al.*, 2007; Julliard *et al.*, 2007; Lacroix *et al.*, 2008; Savigner *et al.*, 2009; Breunig *et al.*, 2010; Tong *et al.*, 2011; Palouzier-Paulignan *et al.*, 2012). Although gating of sensory information at the level of the peripheral receptor cells is widely accepted for most sensory modalities, less is known about modulation at the level of olfactory sensory neurons (Lucero, 2013). Receptors for leptin and insulin are expressed in the olfactory mucosa, and these peptides alter the firing of olfactory sensory neurons in *in vitro* preparations (Baly *et al.*, 2007; Lacroix *et al.*, 2008; Savigner *et al.*, 2009). GnRH has also been shown to reduce electro-olfactogram (EOG) responses in salamanders (Park & Eisthen, 2003). However, the significance of these observations in behaving vertebrates has not been tested. Further, little is known about the molecular mechanisms that underlie changes in OSN sensitivity.

Neuropeptide Y (NPY) has long been recognized as a central modulator of feeding associated behaviours across invertebrate and vertebrate phyla. In *C. elegans* and *Drosophila*, NPY signalling has been implicated in altering chemosensitivity, heightened orexigenic drive, and facilitating presynaptic transmitter release in the OB (de Bono & Bargmann, 1998; Wu *et al.*, 2005; Root *et al.*, 2011; Sengupta, 2013; Ko *et al.*, 2015). In vertebrates, NPY peptide expression in the OE has been reported from teleost fish to mammals (Hansel *et al.*, 2001; Mathieu *et al.*, 2002; Gaikwad *et al.*, 2005). Ectopic application of NPY to OEs isolated from salamanders and rats has been reported to modify the odorant response (Mousley *et al.*, 2006; Negroni *et al.*, 2012).

Using behaving zebrafish, we demonstrate that NPY expression levels in the OE and OB reflect the prevailing energy states, and NPY signalling is necessary and sufficient for gating olfactory responses to food cues. Paracrine signalling by NPY directly modulates the sensitivity of a subset of OSNs to food-associated amino acid cues via a phospholipase C (PLC) dependent mechanism.

## MATERIALS AND METHODS

### Animals

Wild type zebrafish were maintained at 28.5°C on a 14 h light/10 h dark schedule in 10 L and 3 L holding tanks in a recirculating aquarium system (Aquatic Habitats). For studies on zebrafish larvae, the eggs were obtained by setting a cross in 3L breeding tanks (Aquatic Habitats). Eggs were maintained in E3 medium (5 mM NaCl, 0.17 mM KCl, 0.33mM CaCl_2_, 0.33 mM MgSO_4_, 0.00001% methylene blue) in an incubator at 28.5°C till 78 hpf. Unless otherwise indicated, adult fish were fed *ad libitum* twice daily with freshly hatched brine shrimp. Larvae are were fed thrice daily with either the cultured paramecia water or synthetic larval food.The Institutional Animal Ethics Committee approved all the procedures employed in this study.

### Chemicals

Synthetic human neuropeptide Y (NPY), small molecule NPY Y1 receptor inhibitor BIBP-3226 (BIBP), D -(+)-glucose monohydrate (glucose), Trp channel blocker 2-aminoethyl diphenylborinate (2-APB), PLC inhibitor U73122 hydrate (U73122), sodium chloride, paraformaldehyde, sucrose, poly L-lysine, bovine serum albumin (BSA), dimethyl sulfoxide (DMSO), L-lysine, L-histidine monohydrochloride monohydrate, L-tryptophan, L-cysteine hydrochloride, L-alanine, L-phenylalanine, L-valine and L-methionine were purchased from Sigma Aldrich. Gq specific inhibitor GP2A (371780) was purchased from Millipore.

### Intracerebroventricular (icv) and intranasal delivery

Prior to icv delivery, fish were anesthetized following immersion in 2-phenoxyethanol (Sigma-Aldrich, 1:2000). All pharmacological agents were dissolved in 0.9% NaCl or in DMSO depending on solubility. The final concentration of DMSO did not exceed 0.1% for BIBP, 2-APB, GP2A treatments and 0.5% for U73122 treatment. Icv delivery protocol was modified after (Yokogawa *et al.*, 2007). Briefly, 1 µl of solution was delivered directly into the ventricular space using an insulin needle (Becton Dickinson Insulin Syringe U40 – 31G) attached via a catheter to a 10 μl Hamilton microsyringe. After injection, the fish were returned to their tanks and allowed to recover before proceeding for either behavioural recordings or immunohistochemistry. Unless otherwise indicated, all the behavioural and immunofluorescence studies were undertaken 30 mins following injection. The following doses were used for icv delivery of glucose (8000 ng); BIBP (100 pmol); 2-APB (100 µM); GP2A (20 µM); U73122 (1 µM) and NPY peptide (1 pmol).

Intranasal delivery of BIBP-3226 (66.67pmol) or vehicle was carried out by superfusing both the OE by delivering 5µl of the solution via the nares of anesthetized fish for 30 secs using a Hamilton microsyringe. The mouth and gills were covered with a small piece of tissue paper to avoid the solution from spreading. Following the procedure, fish were returned to the tank after 1 minute, allowed to recover from anesthesia for 2 mins and evaluated in the odorant preference assay.

### Amino acid stimulation of the OE for evaluation of OSN activation

Post 30 mins icv delivery of pharmacological agents, fish were transferred into the habitat tank containing 1 liter aquarium water. A mixture of 8 amino acids (Ala, Cys, His, Lys, Met, Phe, Trp, and Val; 0.8 mM each in distilled water or solvent was introduced into these tanks using a syringe. For dose-response experiments, 0 mM, 0.008 mM, 0.08 mM and 0.8 mM concentrations of each of the amino acids was used in the odorant mixture. Following the 8 min exposure, the fish were immediately anesthetized and processed for pERK immunofluorescence.

### Immunofluorescence

Adult fish were anesthetized followed by the removal of skull and immersion in 4% paraformaldehyde (for 20 hours at 4°C) for all experiments, except those involving pERK staining in the OE. In latter experiments, the fixation time was between 14-16 hours in 3% paraformaldehyde at 4°C. The brain and/or the olfactory epithelia were dissected out and transferred to 25% sucrose solution for cryoprotection, embedded in OCT compound (Jung), sectioned in the transverse plane at 10 – 20 μm thickness using a cryostat (CM1950; Leica) and mounted on poly-L-lysine coated slides. Zebrafish larvae (78 hpf) were directly immersed in 4% paraformaldehyde, similarly processed and sectioned in sagittal or transverse planes at 10 μm thickness. The slides containing the sections were stored at −40°C until they were processed for immunofluorescence. The frozen sections were rehydrated and permeabilized using 0.5% Triton-X in phosphate buffer saline (PBST). The sections were treated with 5% heat inactivated goat serum (HIGS) or 5% BSA in PBST for 1 hour and then incubated with the primary antibody overnight at 4°C. The sections were washed using 0.1% PBST and pre-incubated with 5 % HIGS or 5 % BSA in PBST for 1 hour and was incubated with desired secondary antibody for 3 hours at room temperature (RT). Then the sections were washed and mounted with mounting media (70% glycerol, 0.5% N-propyl gallate in 1 M Tris, pH 8.0, DAPI). The following primary antibodies were used in this study – polyclonal anti-NPY antibody (1:2000; Sigma), monoclonal anti-calretinin antibody (1:3000; Swant), monoclonal anti-NPY antibody (1:2000; Sigma), polyclonal anti-G_α/olf_ antibody (1:1000; Santa Cruz), anti-Hu antibody (1:100; Abcam), anti-SV2 antibody (1:50; Developmental Studies Hybridoma Bank), monoclonal anti-phosphorylated ERK 1/2 (1:500; Abcam) and polyclonal anti-TrpC2 (1:50; Abcam). Anti-rabbit Alexa-488 (1:500; Invitrogen) and anti-mouse Alexa-568 (1:500; Invitrogen) were used as secondary antibodies. Sections were observed using either an upright epifluorescence microscope (Axioimager.Z1 with Axiocam HRm camera; Carl Zeiss) or inverted laser scanning confocal microscopes (LSM 710 or LSM 780; Carl Zeiss).

Preadsorption with NPY peptide was carried out to demonstrate the specificity of the anti-NPY antibody (Sigma) used in this study. Briefly, anti-NPY antibody was preadsorbed with synthetic human peptide (N5017; Sigma) at 10^−5^ M for 24 hours. Application of the preadsorbed antibody resulted in total loss of the immunoreactivity from the olfactory bulb sections, confirming the specificity of the antibody (data not shown).

### Whole-mount immunofluorescence

Whole-mount immunohistochemistry was carried out using standard protocols. Briefly, adult zebrafish brain (olfactory nerve and OE intact) were fixed in 4% PFA and post-fixed in Dent’s fixative (4:1 methanol:DMSO) overnight. The brain was then dehydrated in 100% methanol for 30 min. To promote permeabilization, the tissue was subjected to three freeze (−80°C) thaw (RT) cycles in 100% methanol followed by overnight storage at −80°C. The tissue was rehydrated serially with 50% methanol, 15% methanol and PBS for 60 min each at RT. The tissue was permeabilized with proteinase K (10 µg/ml in PBS) for 5 min followed by three washes in PBS. Blocking was carried out for 8 hours at 4°C in 5% BSA, 5% DMSO in 0.5 % Triton X-100. The tissue was then incubated in primary antibody against neuropeptide Y (1:5000) and synaptic vesicle-2 (1:50) for 48 hours at 4°C. Following three washes with PBST (15 min each; 4 °C, the tissue was incubated overnight with secondary antibody Alexa Fluor-488 (1:500) and Alexa Fluor-568 (1:500) at 4°C. The tissue was rinsed three times in PBST (2 hours each; RT) and once in PBT (RT) and finally mounted in 70% glycerol, 0.5% N-propyl gallate in 1 M Tris, pH 8.0, DAPI on a coverslip-bottomed petridish. Images were taken using the inverted laser scanning confocal microscope (LSM 780; Carl Zeiss).

### Quantification of NPY fluorescence intensity and cell count analysis

The intensity of NPY immunoreactivity in the olfactory bulb was measured using Image Pro Plus 7.1 software (Media Cybernetics). Briefly, the precise volume measurements of the bulb were made in the 3D constructor using interactive thresholding. The integrated optical density was then measured in the defined volume of the bulb for NPY and SV2 channels separately. For each treatment, the optical density of the NPY immunoreactivity was normalized with respect to the SV2 immunoreactivity and again with a unit volume defined. NPY immunoreactivity in the soma of the terminal nerve neurons was measured using Image J. Relative NPY fluorescence per terminal nerve neuron in each group was plotted with respect to that of the control group. The intensity of NPY immunoreactivity in the olfactory epithelia was measured using Image J. The total fluorescence was calculated by the formula corrected total fluorescence (CTF) = Integrated density – (Area of the selected ROI * Mean fluorescence of the background readings). The CTF was calculated for both NPY immunoreactivity and DAPI labeling and then the NPY fluorescence was normalized to DAPI fluorescence.

The number of NPY-ir glomeruli in the olfactory bulb and the number of NPY/pERK-ir OSNs in the olfactory epithelia were manually counted. The number of NPY or pERK expressing OSNs were represented as the average number of cells per lamella across all sections covering both epithelia of each individual fish.

### Odorant preference assay in adult zebrafish

In order to habituate the fish (3-5 months old) to the experimental tank (4 × 20 × 17 cm filled with 1400 mL of aquarium water) and the icv injections protocol, a two-day acclimation regime was carried out. On the first day, the fish were anesthetized and injected with saline (0.9 % NaCl) via the icv route. On the second day, a mock injection procedure was followed in which the fish was subjected to the entire exercise as on the first day but no active agent was delivered. During the two-day habituation, the fish were not given any food. The habituated fish were then starved for yet another day and then anaesthetized and injected with either vehicle or BIBP-3226 (100pmol) via the icv route. Similarly, the assay was carried out for the regularly fed fish, starved fish without any injection, and starved fish injected with glucose (8000ng), NPY peptide (1 pmol) or a co-injection of glucose and NPY peptide. Following an interval of 30 mins, the fish were subjected to the odorant preference assay, one at a time, while being video recorded (Sony Handycam DCR SR20) from the side. The olfactory behavioural assay was modified after Koide et al. (2009). Adult zebrafish were assayed for response to a mixture of 8 amino acids (Ala, Cys, His, Lys, Met, Phe, Trp, and Val; 0.1 mM each in distilled water; Sigma). No stimulation was provided for the first 5 mins following which the amino acid mix was introduced in one side of the tank at the rate of 1 ml/min using a syringe pump. Simultaneously, distilled water was delivered at the same rate at the opposite end of the tank. The behaviour of the fish was monitored for a total of 10 mins after the introduction of the odorants. All behavioural assays were performed between 1200h to 1700h.The swimming paths were analyzed from the recorded videos using an automated video tracking system (Ethovision XT version 8.0; Noldus). Preference indices were calculated using the formula: Preference index (PI) = ((Total time spent in the odorant zone) – (Total time spent in the control zone)) / (Total time). The data were evaluated for the period of 5 mins before and after odorant stimulation.

### L-cysteine aversion assay in 78 hpf larvae

The protocol described by (Vitebsky *et al.*, 2005) was employed with modifications. Briefly, 78 hpf zebrafish larvae (n=10-20) were placed in a 35 mm Petri dish containing 5 ml of E3 medium. A circle of 1 cm diameter was drawn on white paper and placed under the petri dish such that the circle was at the center of the dish. The larvae were gently swirled to position them within the perimeter of the circle. L-cysteine (10 ul of 3 × 10^−5^ M) was gently introduced over the surface of the water, approximately at a point above the center of the circle (ensuring minimal vibration of the water surface). The entire procedure was video recorded and the number of larvae that moved out of the area of the circle within 30 secs of addition of L-cysteine was scored. The larvae that remained within the circle were considered as non-responsive. The larvae once used for the assay were not used in further experiments and the entire experiment was carried out at 27-28.5°C.

### Response to alarm pheromone (Schreckstoff)

For testing the alarm response in fish under different energy states, the skin extract was prepared as described earlier (Mathuru *et al.*, 2012). Briefly, several shallow cuts were made on the skin of the zebrafish using a sharp blade. The injured fish was immersed in 2 ml of 20mM Tris-Cl (pH 8.0) for 30 seconds. This process was repeated for 7-10 fish and the extract was collected into the same volume. The 2 ml of crude extract was then centrifuged at 13.2k rpm and the supernatant was collected. A 25% solution of the alarm pheromone was prepared by diluting in distilled water and 500 µl of the latter was used in the behavioural assay.

As stated above in the odorant preference assay, adult zebrafish were singly housed, habituated for 2 days, starved for yet another day, anaesthetized and injected with either saline or glucose (8000ng) via icv route. Following an interval of 30 mins, the fish were subjected to the assay. Control stimulus (500µl of distilled water) was introduced into the tank following the first 5 mins using a plastic dropper. After a period of 10 mins, test stimulus (500 µl of the alarm pheromone extract) was added. The fish were evaluated for their response to the alarm pheromone for 10 minutes. The assay was video recorded (Sony Handycam DCR SR20) and analyzed using an automated video tracking system (Ethovision XT version 8.0; Noldus).

Darting was defined as rapid swimming with increased reversal frequency and swimming velocities at least 2.5 times higher than those recorded in the first 5 mins of observation. Freezing was defined as a period of complete immobility, typically at bottom of the tank for at least 5 seconds.

### NPY Y1 Receptor RNA in situ hybridization

For whole-mount RNA *in situ* hybridization (ISH) experiment adult fish were starved for three days and anesthetized followed by the removal of the dorsal cranium and immersion in 4% paraformaldehyde (for 14-16 hours at 4°C), olfactory rosettes were dissected out, washed in phosphate buffered saline (PBS), and stored in 100% methanol at −20°C overnight. ISH was carried out as described earlier (Akash *et al.*, 2014). Digoxigenin-labeled sense and antisense riboprobes were transcribed *in vitro* using Zebrafish NPY Y1R TOPO pcDNA 3.1 vector template (gift from D. Larhammar) using the following primer pair. Forward: 5’-GACACCAACTCATCCTGCACC-3’; Reverse: 5’-TGGGTACAGTTCATCGCCAC-3’. Probes were purified using BioRad Micro Bio-Spin columns as per manufacturer’s protocol. Samples were sequentially rehydrated to PTW (phosphate buffered saline, 0.1% Tween-20 pH 7.3) and permeabilized in 2% hydrogen peroxide for 35 minutes (min) at room temperature (RT). Samples were further permeabilized by a 40-min proteinase K treatment and post-fixed for 20 min in 4% paraformaldehyde at RT. Prehybridization, probe hybridization and washes were performed as described previously (Akash *et al.*, 2014) except for the inclusion of 5% dextran sulfate in the hybridization buffer. The OE were embedded in 4% low melting point agarose blocks and sectioned (30 µm) using a Vibratome (VT 1200; Leica). The sections were blocked in 1% blocking solution (10 min at RT; Roche Applied Sciences) and then incubated overnight at 4°C with sheep anti-DIG alkaline phosphatase conjugated antibody (1:2,000; Roche Applied Sciences). Sections were washed thrice with PTW buffer and twice with coloration buffer (100 mM Tris-HCl, pH 9.5, 50 mM MgCl_2_, 100 mM NaCl, 0.1% Tween 20, 2 mM Levamisole). DIG-labeled probes were detected using pre-warmed BM purple solution (Roche). The reaction was continuously monitored at 37°C for 30 min to 2 h or overnight at 4°C depending on the strength of the signal. The reaction was stopped by washing three times with PTW buffer. The sections were then mounted in 70% glycerol and were either processed for imaging or stored at 4°C. DIC photomicrographs were taken with an upright microscope (Axioimager Z1, Carl Zeiss).

### Statistical analysis

All the raw measurements were imported to GraphPad Prism 5 (GraphPad Software Inc.) for graphical representation and statistical analysis. In the odorant preference assay, the preference index for each group is represented as boxes and whiskers; the box ranges from the 25^th^ percentile to the 75^th^ percentile and the whiskers represent the minimum and maximum values of the group. The horizontal line in the box denotes the median. The medians of the preference indices, before and after the odorant stimulation, were tested for significant deviation from 0 using Wilcoxon signed-rank test.

The effect of energy status on amino acid sensitivity in the OSNs and the amino acid dose response on animals with different energy states was compared using Two-way ANOVA followed by Bonferroni’s post-hoc test. The confidence limit for statistical significance in all the analyses was p< 0.05.

## RESULTS

### NPY is enriched in the olfactory epithelium and olfactory bulb in zebrafish

Whole mount immunofluorescence analysis of the intact olfactory epithelium (OE) and olfactory bulb (OB) of adult zebrafish with antibodies against NPY and the pre-synaptic marker SV2 (synaptic vesicle transmembrane proteoglycan 2) revealed two sources of NPY in the peripheral olfactory circuitry. First, a subset of the olfactory sensory neurons (OSNs) with NPY fiber projections over the olfactory nerve to the bulb, finally terminating as glomeruli in the OB (Figures 1A, B). The second source of NPY was the terminal nerve (TN). TN soma and their axonal projections innervating the OE as well as the extrabulbar telencephalic regions, showed extensive NPY expression (Figure 1A).

**Figure 1.**
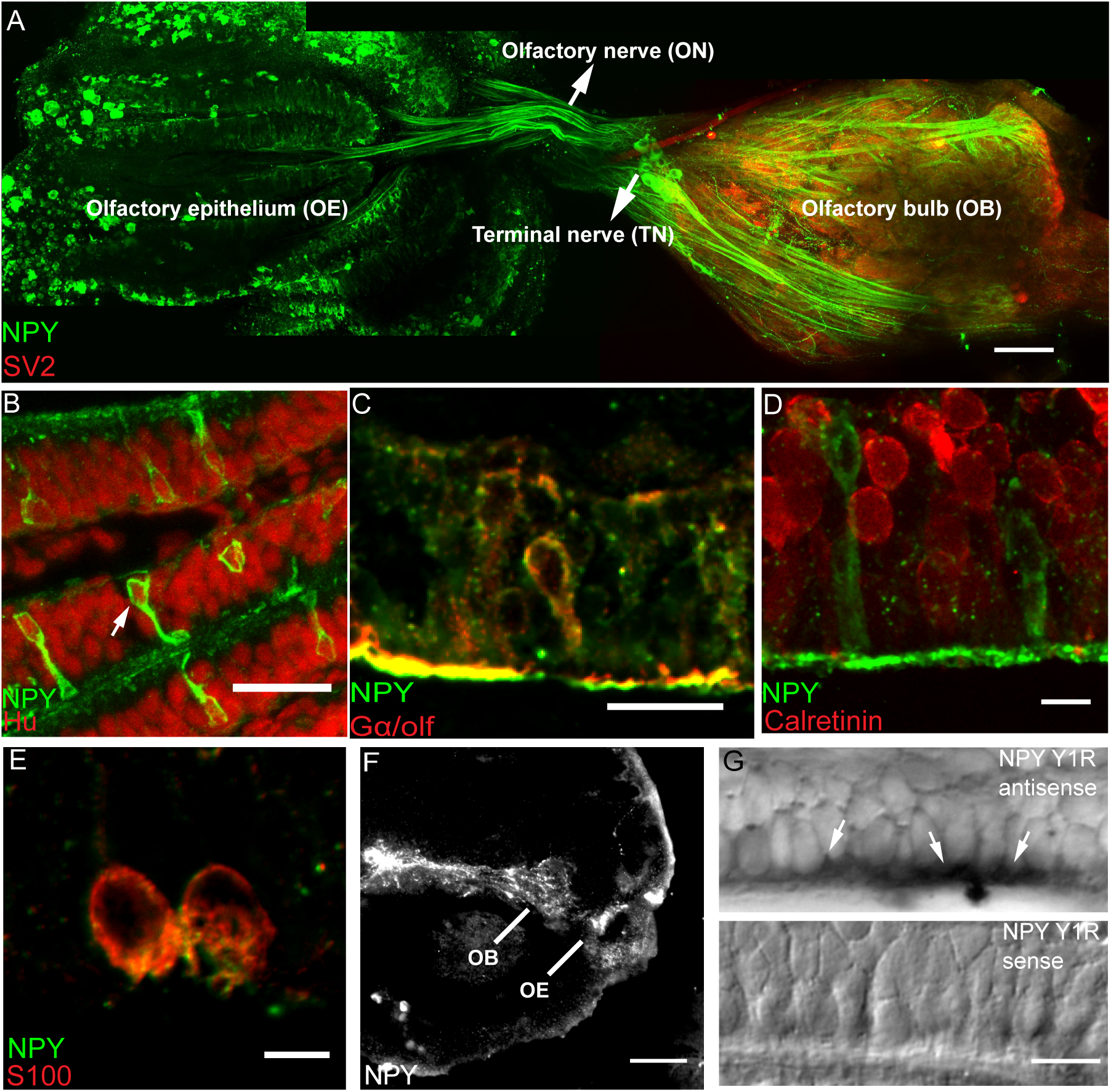
NPY and NPY Y1 receptor expression in the zebrafish olfactory system. **(A)** Whole mount immunofluorescence of adult zebrafish OE and OB. NPY(green) is expressed in the OE, OB, ON (arrow) and TN neurons (arrow) of adult zebrafish. NPY immunoreactive (NPY-ir) axon terminals synapse on glomeruli marked with SV2 (red). **(B)** Transverse section across olfactory epithelium showing the colocalisation (arrow) of NPY-ir OSNs (green) with Hu (red). **(C,D,E)** Transverse sections across olfactory epithelium showing colocalisation of NPY (green) with G_α/olf_ (red, **C**) and S100 (red, **E**) but not with Calretinin (red, **D**). **(F)** Saggital section showing NPY immunoreactivity in the OE and OB of 78 hpf zebrafish larvae. **(G)** Representative RNA *in situ* hybridisation using NPY Y1 receptor mRNA antisense (upper panel) and sense (lower panel) probes showing expression in adult zebrafish OSNs (arrows). [Scale bars: **A**, 200 μm; **B**,**D**,**G** 10 μm; **F**, 50 μm; **C**,**E**, 5 μm]

Double immunostaining with NPY and neuronal marker Hu (Figure 1B) (Iqbal & Byrd-Jacobs, 2010) confirmed the neuronal identity of the NPY expressing cells. Co-localization with OSN-subtype markers revealed that NPY expression was largely limited to the crypt (co-localization with S100) and G_olf_ positive ciliated OSNs. However, it was excluded from calretinin-expressing microvillar and ciliated OSN populations (Figures 1C-E).

Expression of NPY in the olfactory anlage of larval zebrafish was tested using immunofluorescence in 78 hpf zebrafish. Consistent with the observations in adult fish, sagittal sections of 78 hpf zebrafish larvae showed NPY expression in both the OE and OB (Figure 1F).

NPY signalling via the NPY Y1 receptor (Y1R) has been implicated in regulating food intake in multiple species (Mercer *et al.*, 2011; Nguyen *et al.*, 2012; Yokobori *et al.*, 2012). Further, Y1R mediates olfactory processing in rodents (Negroni *et al.*, 2012). mRNA *in situ* hybridization using antisense probe against Y1R mRNA showed intense signal in subpopulations of OSNs (Figure 1G upper panel) while the sense probe controls did not produce any reaction (Figure 1G lower panel).

These data demonstrate that NPY and the NPY Y1 receptor is expressed in the peripheral olfactory circuitry of zebrafish.

### Energy status-dependent expression of NPY in the olfactory system

NPY promotes hunger and increases food intake across vertebrate phyla, including zebrafish and its expression is elevated in multiple neuronal regions upon fasting (Clark *et al.*, 1987; Marks *et al.*, 1992; Volkoff *et al.*, 2005; Yokobori *et al.*, 2012). Fed (free feeding on an excess of food 30 mins prior to sacrifice; mock icv injection), starved (for 3 days; mock icv injection) and adult zebrafish injected with glucose (via icv route) were compared for NPY expression in the olfactory epithelia and olfactory bulb. Glucose icv treatment allowed isolation of central regulation from peripheral homeostatic inputs. The latter treatment also reduced the variability in free feeding fish that may arise from individual proclivities. Total fluorescence intensity of the NPY signal and the number of NPY expressing OSNs in the olfactory mucosa increased upon starvation in comparison to either well fed or glucose injected animals (Figure 2A-C, Figure 2J, p= 0.0044 and Figure 2K, p= 0.0065). Similarly, both the NPY signal intensity in the OB and the number of NPY-positive glomeruli also increased upon starvation (Figure 2D-F, Figure 2L, p<0.0001 and Figure 2M, p= 0.0059). NPY fluorescence in the olfactory bulb comprises of signals from both the OSN projections onto glomeruli and the terminal nerve. Independent evaluation of the terminal nerve showed an increase in NPY immunofluorescence per TN soma and also in the number of NPY expressing TN cell bodies in starved zebrafish compared either to fed or glucose injected fish (Figure 2G-I, Figure 2N, p<0.0001 and Figure 2O, p<0.0001).

**Figure 2.**
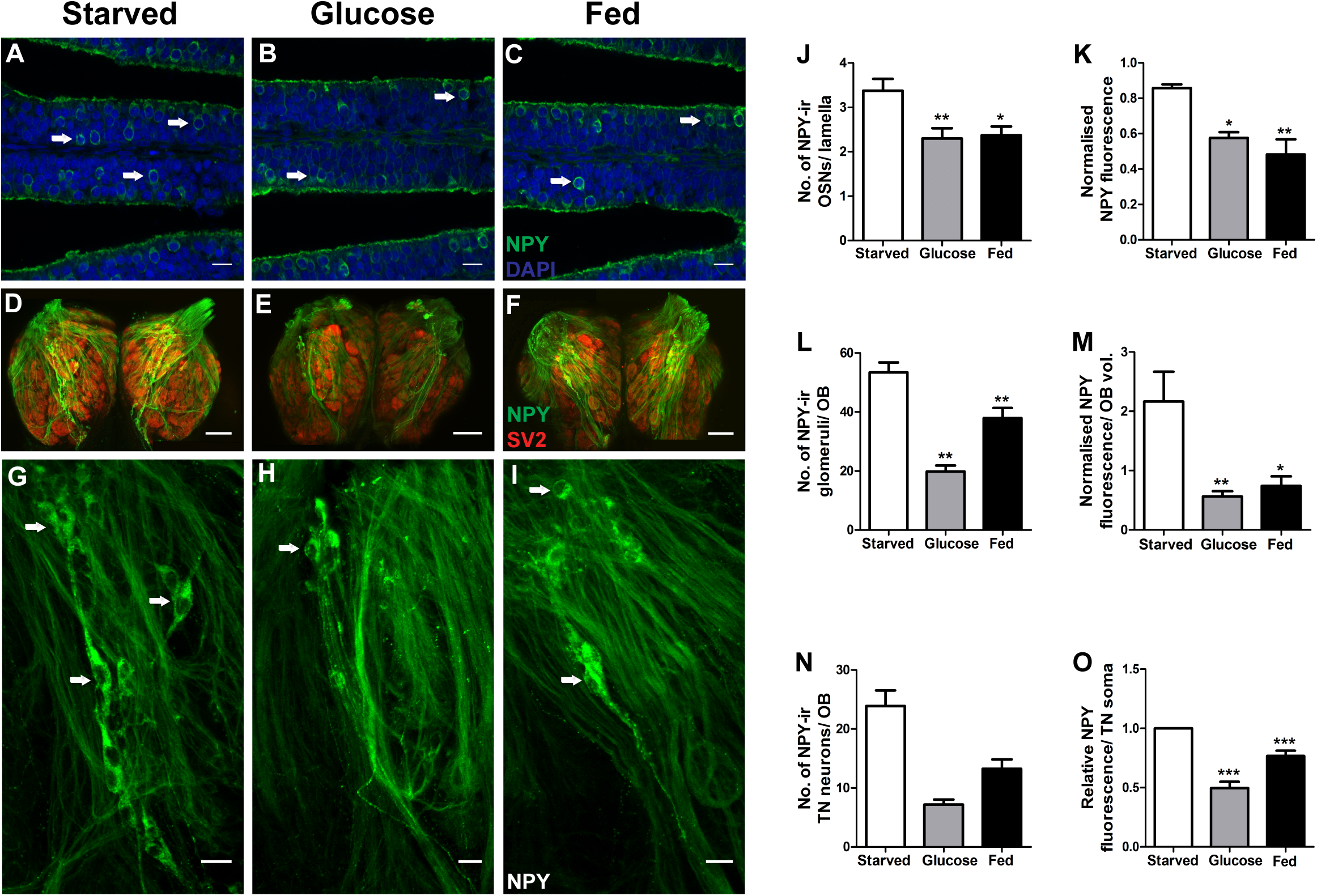
NPY expression in the olfactory system is sensitive to nutritional states. **(A,B,C)** Representative transverse sections showing NPY immunoreactivity (green) and DAPI (blue) in the OSNs (arrows) of the olfactory organ for different feeding states (Starved, Glucose and Fed). **(D,E,F)** Whole mount immunofluorescence of olfactory bulb showing NPY (green) and SV-2 (red). **(G,H,I)** NPY-ir in the terminal nerve neurons (arrows) in the olfactory bulb for different feeding states. **(J)** Number of NPY-ir OSNs per lamella across all 20µm sections for both the OEs under different feeding states (N=3). **(K)** Normalised NPY fluorescence in the whole olfactory organ for different feeding states (N=3). **(L)** The number of NPY-ir glomeruli in the whole olfactory bulb for different feeding states (N=6). **(M)** Normalised NPY fluorescence in the whole olfactory bulb for different feeding states (N=5). **(N)** Number of NPY-ir terminal nerve (TN) neurons per each olfactory bulb (N=6) and **(O)** The relative intensity of NPY-ir TN neurons in the whole olfactory bulb for different feeding states (N=6). Data are shown as mean +/-SEM (*, p<0.5, **, p< 0.01, ***, p < 0.001; One – way ANOVA followed by Bonferroni post hoc test analysis for significance in comparison with the starved group). [Scale bars: **A-C**,10 µm: **D-I**, 20 µm].

These results indicate that NPY expression levels in the olfactory epithelia and bulb reflect the prevailing energy status in the brain.

### Nutritional state-dependent olfactory perception of amino acids is mediated by NPY Y1 receptor signalling

Energy state-dependent NPY expression and availability of NPY-receptors in the peripheral olfactory circuitry of zebrafish could be indicative of a role for NPY signalling in nutritional state dependent olfactory processing. We evaluated olfactory responses of fed and starved adult zebrafish in an odorant preference assay using a mixture of amino acids, a major food-associated attractive odorant cue in fish (Figure 3A). The percent time spent in the odorant half by starved animals was more in the post-odorant period compared to the pre-odorant stimulation period and was reflected in the statistically significant increase in the preference index (PI) for the baited side (Figure 3C, p= 0.0156). However, fed animals displayed no difference in PI between the pre- and post-stimulation periods (Figure 3B).

**Figure 3.**
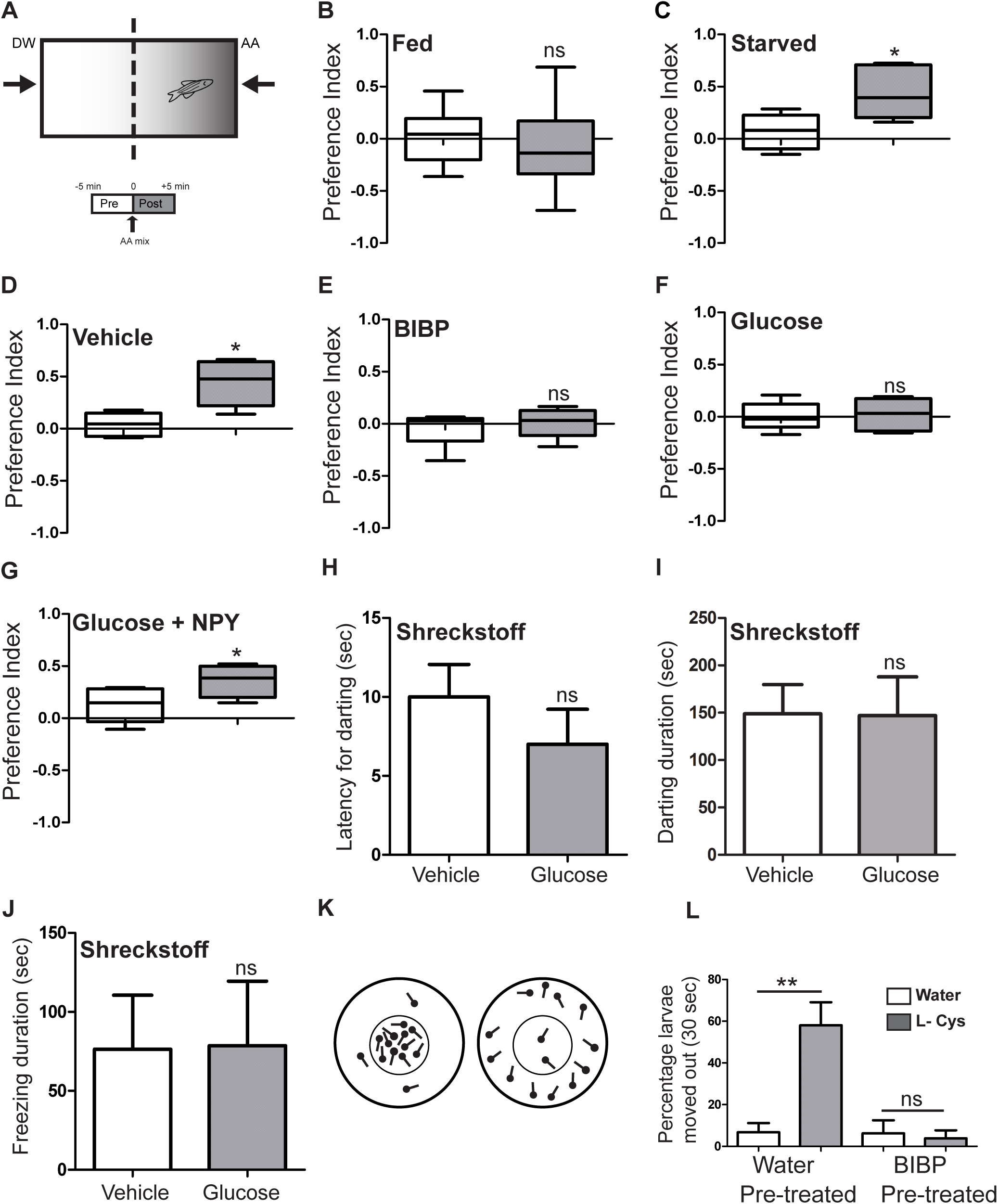
NPY signalling is necessary and sufficient for olfactory responses to food-associated amino acid cues. **(A)** Schematic of the odorant preference assay in adult fish using a mixture of amino acids as an attractive odorant cue (see methods for explanation) **(B,C)** Fraction of time spent in the odorant zone as indicated by the preference index in a fed and starved fish (N=6), respectively. The preference index before and after addition of amino acids is tested for significant deviation from 0 using Wilcoxon signed-rank test (*, p< 0.05; ns, no significant difference from 0). **(D-G)** Preference index of starved fish injected via icv with either Vehicle, BIBP, Glucose or co-injected glucose along with NPY peptide (N=6; *, p<0.05 in both D and G; ns, no significant difference from 0). **(H-J)** Starved fish injected with either vehicle (N=10) or glucose (N=7) showed no significant difference (ns) in their response to the alarm cue, Schreckstoff. Data are shown as mean +/-SEM; unpaired t-test was used to test for statistical significance. **(K)** Schematic of the aversion assay in the 78 hpf zebrafish larvae using L-Cysteine as an aversive cue (see methods for explanation). **(L)** 78 hpf larvae pre-treated with water show aversion to L-Cysteine (N=6; **, p< 0.001) while larvae pre-treated with BIBP show no significant (ns) difference between the L-Cysteine treated and control fish. Data are shown as mean +/-SEM; One-way ANOVA followed by Bonferroni’s post-hoc test was used to test for statistical significance.

To test the role of NPY signalling in olfaction, we evaluated odorant preference behaviour of starved fish (high NPY expression in OE and OB) while abrogating NPY signalling through the NPY Y1 receptor (Y1R; the major receptor associated with NPY-dependent appetitive behaviours (Yokobori *et al.*, 2012)). The PIs for animals receiving Y1R antagonist BIBP-3226 (BIBP) via the icv route showed no difference between pre- and post-amino acid stimulation periods (Figure 3E). However, fish receiving the control treatment (saline via icv route) displayed an increased preference for the baited side following odorant stimulation (Figure 3D, p= 0.0313). Analysis of locomotor behaviour revealed no differences between BIBP and control animals (data not shown).

As NPY expression in the OE and OB was reduced in fed and glucose administered animals compared to starved animals (Figure 2), we tested whether NPY-signalling was sufficient to restore olfactory sensitivity to amino acids in energy-rich conditions. Starved fish receiving glucose via the icv route (shown previously to reduce endogenous NPY expression) showed no difference in PI before and after amino acid baiting (Figure 3F). However, administration of exogenous NPY to glucose-injected animals was sufficient to restore odorant preference (Figure 3G, p= 0.0313).

To test whether NPY signalling modulates nutritional state-dependent olfactory sensitivity specifically for food-associated cues, we evaluated the response to the alarm cue, Schreckstoff. The skin of conspecifics produce Schreckstoff, which is sensed by the crypt subtype of OSNs and robustly triggers a sequence of behaviours that include vigorous darting movements followed by freezing (Mathuru *et al.*, 2012). We compared the response of Schreckstoff between fed and starved adult zebrafish. Schreckstoff application initiated strong behavioural responses, but no differences were detectable between the two nutritional states. The duration of both darting movements (Figure 3I; ns) or immobility (Figure 3J; ns) were comparable between starved and fed animals. As vigorous darting preceded the freezing behaviour, we evaluated the latency to display the darting behaviour as a measure of sensitivity to the alarm cue. However, the latency for darting was also similar between fed and starved zebrafish (Figure 3H; ns).

These experiments suggest that endogenous NPY signalling via the NPY Y1 receptor is subject to regulation by the brain energy status and is sufficient to specifically modulate the olfactory sensitivity to food-associated amino acid cues.

### NPY signalling modulates olfactory responses to amino acids in larval zebrafish

Reduction in NPY signalling has been associated with the attenuation of the feeding drive. Thus, the use of Y1R antagonist and food-associated attractive odorant cues (amino acid mixture) in the odorant preference assay may potentially confound an olfactory response with a generalized loss of the feeding drive. To exclude this possibility, we employed an amino acid –dependent aversive assay. 78 hpf zebrafish were stimulated with L-Cysteine, an aversive olfactory signal at high concentrations (Vitebsky *et al.*, 2005), and the escape response monitored (Figure 3K). While a large proportion of larvae responded to L-cysteine (Figure 3I; p<0.05), pre-treatment with BIBP abolished the aversive response, which was indistinguishable from the control groups (Figure 3L; ns). This assay, using an amino acid-based aversive cue, demonstrates that NPY-mediated behavioural responses are not an artifact of reduced feeding drive. Rather, NPY signalling modulates amino acid sensing.

The L-cysteine aversion assay in larvae also addressed a second potential confound. While the adult odorant preference assay is largely specific to the olfactory system (Koide *et al.*, 2009), it has also been reported that the fish gustatory system can sense amino acids (Oike *et al.*, 2007). However, as the gustatory system is not functional at the developmental time of 78 hpf (Lindsay & Vogt, 2004), the L-cysteine aversion assay at this developmental stage indicates that the olfactory system is the sole sensory modality involved. Collectively, the larval L-cysteine aversion assay implicates NPY signalling in the olfactory processing of amino acid cues.

### Peripheral modulation of olfactory behaviour by NPY

The icv mode of delivery of NPY signalling antagonists in the previous experiments does not allow discrimination between central and peripheral contributions. In order to isolate the role of peripheral modulation, we developed a nasal delivery protocol selectively targeting the peripheral olfactory circuitry. Olfactory epithelia of starved, adult zebrafish were bilaterally bathed with BIBP for 30 secs, introduced via the naris and tested in the odorant preference assay. While the BIBP treated fish were agnostic to the amino acid cues, the control animals were attracted to the amino acid baited side (Figures 4A,B).

**Figure 4.**
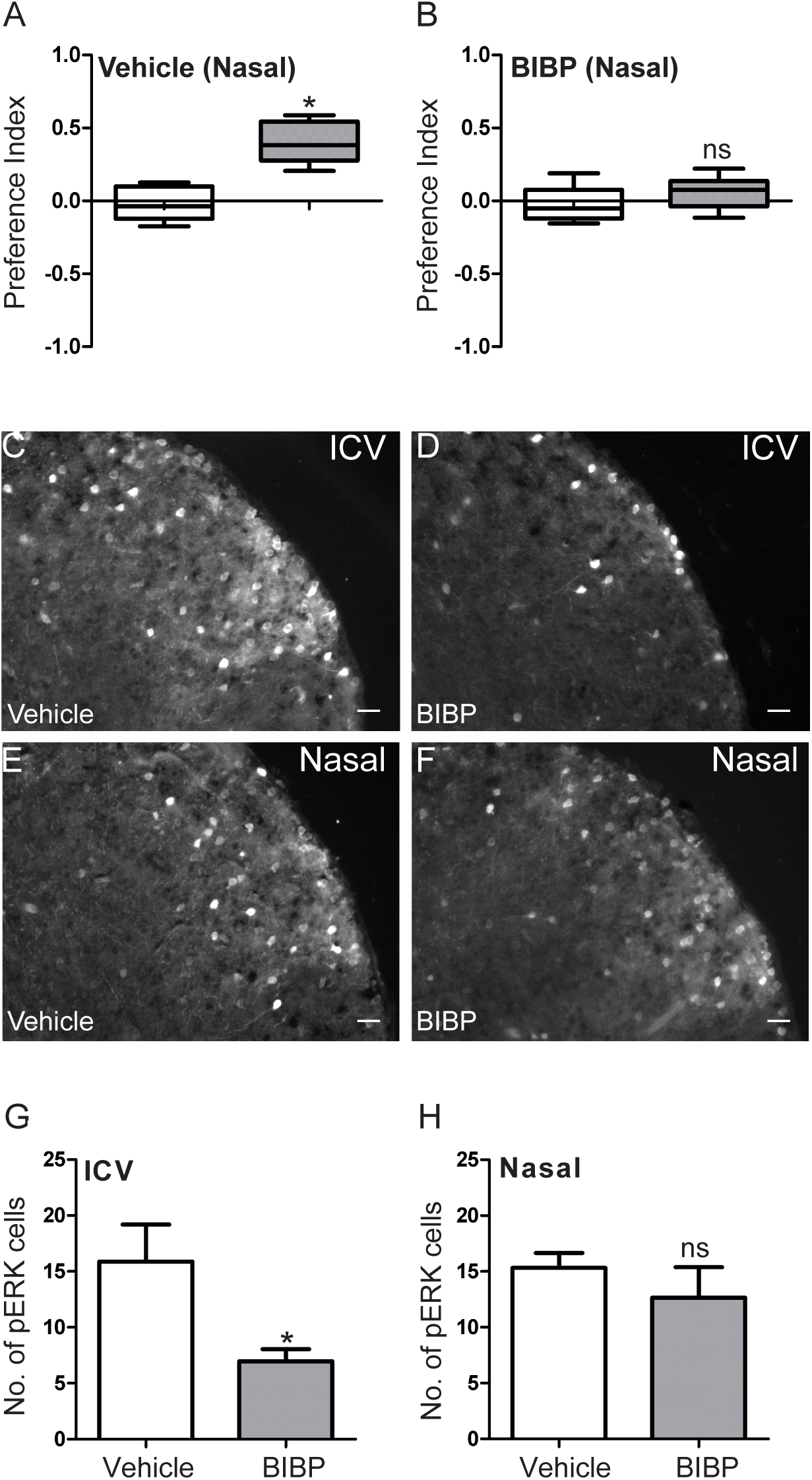
NPY signalling modulates olfactory sensitivity at the periphery. **(A,B)** Preference index of starved fish in the odorant preference assay treated either with vehicle or BIBP via the nasal route (N=6). The preference index before and after addition of amino acids was tested for significant deviation from 0 using Wilcoxon signed-rank test (*, p< 0.05; ns, no significant difference from 0). **(C-F)** Representative micrographs of a Dl region from transverse sections across the telencephalon showing pERK immunoreactivity in starved fish pretreated with vehicle/BIBP via icv (**C,D**) or nasal route **(E,F)**. **(G,H)** The number of pERK-ir cells in Dl region in starved fish injected with either vehicle or BIBP via icv route (N=6; G) or via nasal route (N=6; H). Data are shown as mean +/-SEM; unpaired t-test was used to test for statistical significance. *, p< 0.05; ns, no statistically significant difference. [Scale bars: **C-F**, 25 μm].

Phosphorylated ERK1/2 (pERK) is a sensitive marker of neuronal activation (Hussain *et al.*, 2013; Randlett *et al.*, 2015). The dorsolateral region of the telencephalon (Dl) produce spontaneous neural activity detectable by pERK immunoreactivity. We used this observation to compare the influence of BIBP on the zebrafish telencephalon between the icv and nasal delivery routes. Compared to control treatment, BIBP administered to starved fish via the icv route consistently resulted in a decrease in pERK-positive cells in the dorsolateral region of the telencephalon (Dl) (Figure 4C,D,G, p= 0.0280), suggesting a role for NPY signalling in promoting activity in this region. However, nasal lavage of BIBP, under the same conditions that abrogate odor preference in the behavioural test, failed to decrease ERK activation in the Dl neurons (Figure 4E,F,H, ns). These experiments suggest that the conditions optimized for the nasal delivery of BIBP led to selective inactivation of NPY signalling that did not extend to telencephalic regions.

In summary, the data from nasal delivery of the NPY Y1 receptor antagonist implicate a significant contribution of NPY-signalling in peripheral modulation of olfactory responses to amino acid cues. The nasal delivery experiments implicate regions upstream of the telencephalon, namely the OE and/or the OB as the site/s of action of NPY modulation. This observation led us to directly evaluate the response of OSNs, as described in the next section.

### Peripheral modulation of amino acid sensitivity of OSNs by NPY signalling

The indication that NPY may act in the periphery to modulate olfaction prompted us to investigate the response of the OSNs to amino acids under different conditions. The microvillar subtype of OSNs employ vomeronasal type 2 receptors (V2Rs) to mediate amino acid sensing in teleost fish and amphibia, like *Xenopus* (Koide *et al.*, 2009; Sansone *et al.*, 2014). Unlike classical OSN signal transduction, V2R signalling involves phospholipase C activity and opening of the diacylglycerol (DAG) -gated TrpC2 channels resulting in OSN activation (Lucas *et al.*, 2003; Sato *et al.*, 2005; Koide *et al.*, 2009; Sansone *et al.*, 2014). To test if amino acid mediated activation of OSNs in zebrafish was limited to the microvillar OSNs, we directly evaluated OSN activation by monitoring ERK1/2 phosphorylation (pERK). We found that OSNs activated by amino acid stimulation, as revealed by ERK activity, exclusively co-localized with TrpC expressing OSNs (Figure 5A-C).

**Figure 5.**
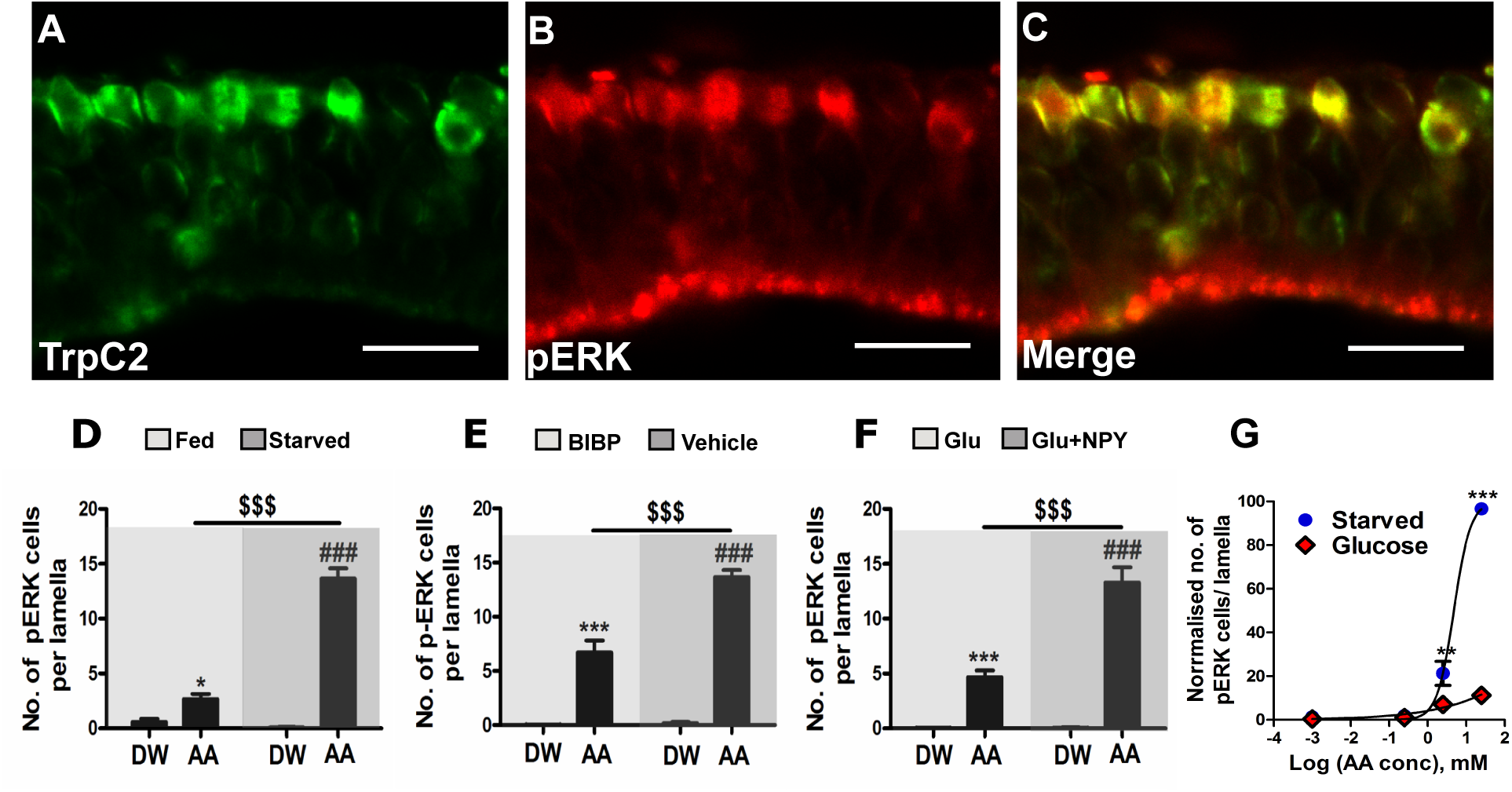
NPY signalling increases the sensitivity of microvillar OSNs to amino acids. **(A-C)** Representative transverse sections showing the immunoreactivity of TrpC2 (green), pERK (red) and their colocalisation (yellow) in the OSNs. **(D)** Effect of starvation on amino acid sensitivity in adult zebrafish. Adult zebrafish were either regularly fed (light gray) or starved (dark gray) for 3 days and were exposed to either a mixture of amino acids (AA) or distilled water (DW). The values are means +/-SEM, n= 6. Post –hoc analysis of significant odorant × energy status interactions in Two-way ANOVA : \**p*< 0.05, Fed (AA) versus Fed (DW); ^###^*p*< 0.001, Starved (AA) versus Starved (DW); ^$$$^*p*< 0.001, Starved (AA) versus Fed (AA). **(E)** Effect of blocking NPY Y1R signalling on amino acid sensitivity in adult zebrafish. Adult zebrafish were starved for 3 days, injected with either BIBP (light gray) or Vehicle (dark gray) and were exposed to either a mixture of amino acids (AA) or distilled water (DW). The values are means +/-SEM, n= 6. Post –hoc analysis of significant odorant × NPY signalling interactions in Two-way ANOVA : \*\*\**p*< 0.001, BIBP (AA) versus BIBP (DW); ^###^*p*< 0.001, Vehicle (AA) versus Vehicle (DW); ^$$$^*p*< 0.001, Vehicle (AA) versus BIBP (AA). **(F)** Effect of energy status on amino acid sensitivity in adult zebrafish. Adult zebrafish were starved for 3 days, injected with either Glucose (Glu; light gray) or co-injected NPY peptide with glucose (Glu + NPY; dark gray) and were exposed to either a mixture of amino acids (AA) or distilled water (DW). The values are means +/-SEM, n= 6. Post – hoc analysis of significant odorant × energy status interactions in Two-way ANOVA : \*\*\**p*< 0.001, Glu (AA) versus Glu (DW); ^###^*p*< 0.001, Glu + NPY (AA) versus Glu + NPY (DW); ^$$$^*p*< 0.001, Gluc + NPY (AA) versus Glu (AA). In D-F, light grey shading indicates conditions of reduced NPY signalling while dark grey shading indicates elevated NPY signalling. **(G)** Normalized response of OSNs to different concentrations of amino acids (log scale) evaluated by counting the number of pERK-ir OSNs in starved fish injected with glucose (red) and vehicle (blue) (N=3; error bars represent +/-SEM; individual p-values obtained from 2-way ANOVA followed by Bonferroni’s post-hoc test; **, p<0.01; ***, p<0.001). [Scale bars: **A-C**, 10 μm].

We next tested if prevailing energy states directly modulated OSN activation by amino acids. Fed and starved zebrafish were stimulated with amino acid or distilled water and OSN activation was evaluated by scoring for pERK signal. A small increase in the number of activated OSNs was observed in fed fish stimulated with amino acids compared to controls (Figure 5D). However, a significantly larger response was seen upon amino acid stimulation of starved zebrafish suggesting increased OSN sensitivity to odorants in these animals (Figure 5D). Two-way ANOVA revealed a statistically significant interaction between the high NPY signalling (starvation) and V2R activation (amino acid stimulation) denoted as the [NPY × AA] interaction (Figure 5D; F_(1,20)_=112.2, p< 0.0001). Post-hoc analysis revealed that while amino acid stimulation had a small effect (increased OSN activation in fed fish where endogenous NPY signalling is reduced compared to the controls), high NPY signalling alone failed to activate OSNs in the absence of odorant (no difference between control treatments of fed and starved fish).

To directly test the role of NPY in OSN activation, we used the NPY Y1R antagonist BIBP administered via the icv route to block NPY signalling in starved zebrafish. Compared to distilled water, amino acid stimulation caused a large increase in OSN activation in starved control animals (Figure 5E). However, the OSN activation was significantly attenuated in BIBP pre-treated starved animals compared to those receiving the vehicle, implicating NPY signalling in tuning OSN sensitivity (Figure 5E). As in the previous experiment, comparing fed and starved fish, two-way ANOVA revealed a statistically significant [NPY × AA] interaction (Figure 5E; F_(1,20)_= 28.06, p< 0.0001). While post-hoc analysis showed an effect for amino acid stimulation (increase in OSN activation by amino acid compared to distilled water stimulation in BIBP treated fish), increase in NPY signalling alone was not enough to change OSN activation above baseline (no difference between distilled water stimulated BIBP and vehicle administered fish).

To test if NPY signalling was sufficient to restore OSN sensitivity to amino acid stimulation, we evaluated OSN activity (pERK-ir) in fish where endogenous NPY signalling is suppressed by administration of glucose via the icv route followed by addition of exogenous NPY peptide. Compared to distilled water, amino acid stimulation resulted in an increase in the number of OSNs activated in glucose-injected fish (Figure 5F). However, this change was significantly smaller than the large increase in OSN activation seen upon amino acid stimulation of glucose-injected fish that have also received exogenous NPY (Figure 5F). These data implicate NPY signalling to be sufficient in restoring OSN sensitivity to amino acids. Again two-way ANOVA revealed a statistically significant [NPY × AA] interaction (Figure 5F; F_(1,20)_= 31.71, p< 0.0001). As in the last two experiments, amino acid stimulation did have a statistically significant effect (amino acid increased OSN activation compared to distilled water in glucose-injected animals) but this was significantly smaller than the combined effect of high NPY signalling and amino acid stimulation.

In all the conditions tested, two-way ANOVA revealed a consistent statistically significant [NPY × AA] interaction. This strongly suggests a cooperative effect between NPY signalling and amino acid triggered signal transduction (via the V2Rs), which is not merely additive. This conclusion is further strengthened by the observation that artificially increased NPY signalling by injecting starved fish (high endogenous NPY expression) with additional NPY peptide fails to show any OSN activation in the absence of amino acid stimulation (data not shown).

These experiments suggest that increase in NPY in the peripheral olfactory system of energy-deficient animals may lower the threshold for OSN activation by food-related odorants (amino acids), while the converse holds for the energy-rich state. To test the prediction that nutritional states alter OSN sensitivity to amino acid cues, we compared the extent of OSN activation in glucose administered (via icv) and starved fish (saline via icv) in response to varying concentrations of the amino acid cues. Two-way ANOVA revealed a statistically significant difference between the two dose-response curves (Figure 5G; F_(1,16)_= 273.27, p< 0.0001) with the starved fish showing a sharp increase in sensitivity to amino acid cues compared to glucose treated animals. This result supports the conclusion that heightened NPY signalling in starved animals increases the sensitivity of OSNs to amino acid cues.

Taken together these experiments underscore the dynamic modulation of the OSN activation threshold in different energy states and implicate NPY signalling acting cooperatively with amino acid signal transduction in tuning OSN sensitivity to food cues.

### NPY and amino acid signalling converges on PLC to activate OSNs

Our experiments suggest that NPY signalling facilitates olfactory perception of food-associated amino acids, which is largely limited to microvillar OSNs that employ a V2R-PLC-TrpC2 signalling cascade. Attenuation of OSN activation by blocking TrpC2 activity would be confirmatory but there are no specific TrpC2 inhibitors currently available. We therefore used 2-APB, a general Trp channel blocker (not specific for TrpC2), that has been shown to block OSN responses in the vomeronasal organ in rodents (Lucas *et al.*, 2003; Zhang *et al.*, 2010) and amino acid responses in the OE of *Xenopus* (Sansone *et al.*, 2014). Pretreatment of zebrafish with 2-APB administered via icv route significantly attenuated OSN activation by amino acids in starved zebrafish (Figure 6A, p<0.0001).

**Figure 6.**
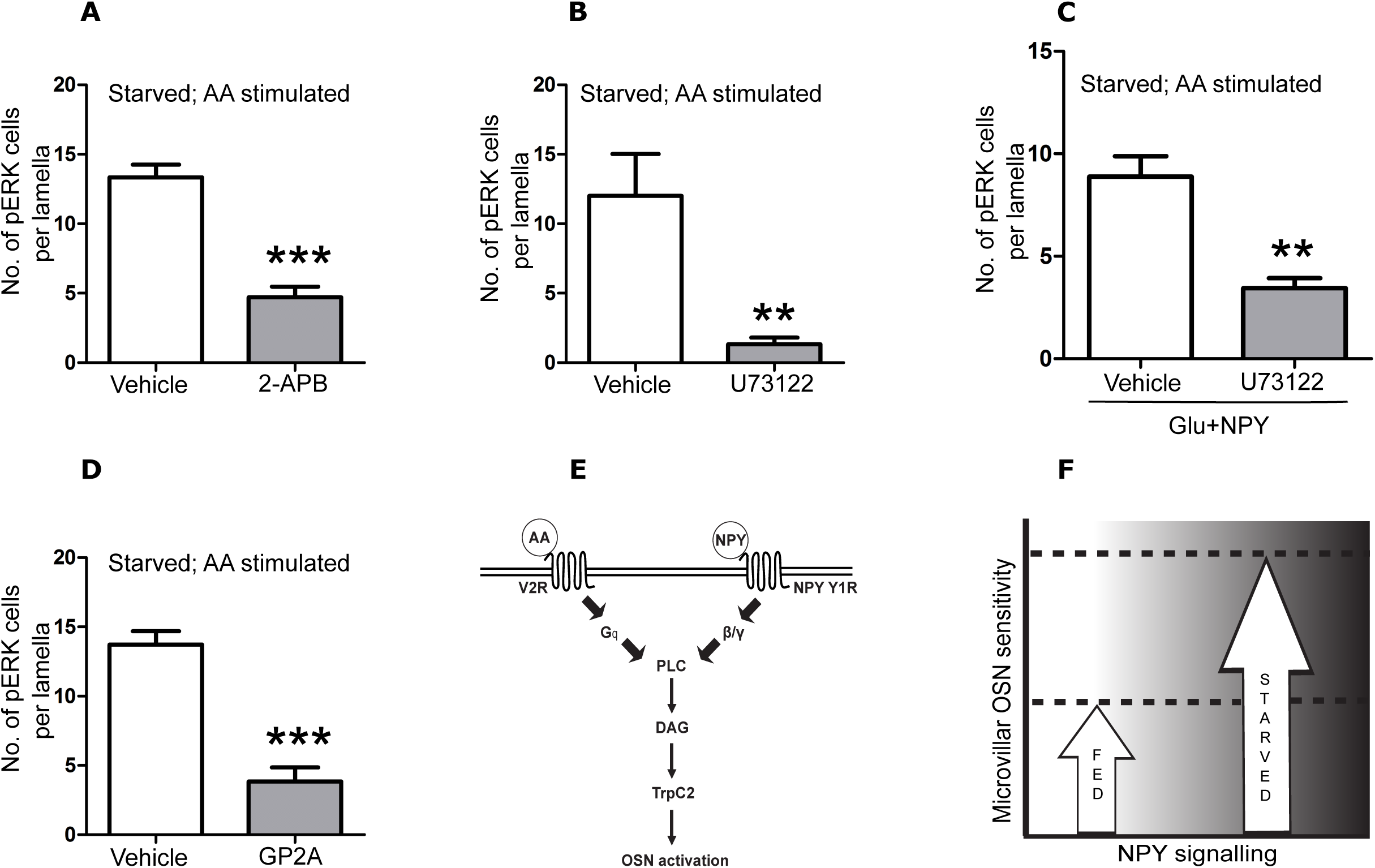
Amino acid and NPY signalling converge on PLC activity to modulate OSN sensitivity. **(A)** The number of pERK –ir OSNs in starved fish injected with either vehicle or 2-APB (Trp channel blocker) via icv route and exposed to a mixture of amino acids (N=6; error bars represent +/-SEM; ***, p< 0.0001). **(B)** The number of pERK –ir OSNs in starved fish injected with either vehicle or U73122 (selective PLC inhibitor) via icv route and exposed to a mixture of amino acids (N=6; **, p< 0.001). **(C)** The number of pERK –ir OSNs in starved fish pre-injected with Glu + NPY followed by administration of vehicle or U73122 via icv route, exposed to a mixture of amino acids (N=6; **, p< 0.01). **(D)** The number of pERK –ir OSNs in starved fish injected with either vehicle or GP2A (selective G_αq_ signalling inhibitor) via icv route and exposed to a mixture of amino acids (N=6; ***, p< 0.0001). Data are shown as mean +/-SEM and means were compared using unpaired t-test. **(E)** Schematic showing the convergence of NPY Y1 receptor (NPY Y1R) signalling and odorant receptor for amino acids (V2R) signalling on phospholipase C mediated activation of microvillar OSN. **(F)** Increased NPY signalling (starved animals) results in a sharp increase in the sensitivity of microvillar OSNs to food-associated amino acid odorant cues. In energy-rich conditions (fed animals), the baseline NPY signalling is low and the sensitivity of the microvillar OSNs to amino acids is reduced. AA – amino acid mixture; PLC – phospholipase C; DAG – diacylglycerol; TrpC2 – transient receptor potential channel C2.

As phospholipase C is central to V2R signal transduction, inhibition of PLC should attenuate OSN activation by amino acids. Indeed, pretreatment with a generic PLC inhibitor, U73122 (introduced via the icv route), resulted in a strong reduction in the number OSNs showing amino acid induced pERK expression compared to animal pretreated with the vehicle control (Figure 6B, p= 0.0058). This suggests a role for PLC signalling in processing amino acid cues in the olfactory epithelia.

It has been shown earlier that exogenous NPY is sufficient to restore both the behavioural response to amino acids, and activation of OSNs by amino acids under conditions where endogenous NPY is reduced by glucose administration (Figures 3G and 5F). We tested if the restoration of OSN activation by exogenous NPY was dependent on PLC activity. Pretreatment with the PLC inhibitor prevented the increase in pERK positive OSNs upon amino acid stimulation even in the presence of exogenous NPY peptide. In contrast, the vehicle treated animals showed the expected increase in OSN activation (Figure 6C, p=0.0014). These results implicate PLC signalling in NPY-mediated lowering of the amino acid sensitivity threshold of OSNs. NPY receptor mediated activation of PLC has been documented previously and is thought to involve G_β/γ_ signalling (Koch *et al.*, 1994; Selbie *et al.*, 1995; Selbie *et al.*, 1997).

We have shown earlier that NPY receptor and amino acid stimulated V2R signalling have a cooperative effect of OSN activation (Figure 5D-F) and involve PLC activity (Figure 6B, C above). Interestingly, there is a significant body of literature implicating synergistic activation of PLCβ isoforms in GPCR signalling crosstalk (Zhu & Birnbaumer, 1996; Selbie *et al.*, 1997). Cooperative activation of PLCβ necessitates simultaneous occupancy of the G_β/γ_ and G_αq_ binding sites on PLCβ. NPY receptor signalling via G_β/γ_ has been implicated in potentiation of adrenergic and purinergic signalling responses via synergistic activation of PLC (Philip *et al.*, 2010). Assuming a similar cooperative mechanism is operational in zebrafish OSNs where our data show convergent signalling by NPY and amino acids onto PLC, V2R activation is expected to provide the G_αq_ component. We used a G_αq_ specific inhibitory peptide, GP2A, to test the involvement of G_αq_ in amino acid odorant transduction. Starved animals (high endogenous NPY expression) were pretreated with GP2A or vehicle control and stimulated with amino acid mixture. While amino acid stimulation caused a large increase in OSN activation in control fish, GP2A treatment significantly attenuated amino acid activation of OSNs (Figure 6D, p<0.0001). The residual number of activated OSNs was comparable to that seen on amino acid stimulation of BIBP- or glucose-injected animals.

These data implicated PLC signalling as a point of convergence of NPY Y1R and amino acid-sensing V2R signalling. Further, the possibility of cooperative activation of PLC by the two signalling cascades as the biochemical basis of cooperativity observed at the levels of OSN activation is strengthened by the identification of a critical role of G_αq_ signalling in amino acid signal transduction in zebrafish OSNs.

## DISCUSSION

We have used zebrafish to study the modulation of olfactory sensitivity to food-related cues by NPY signalling in response to the prevailing nutritional state. As NPY expression in the peripheral olfactory circuitry was tuned to the nutritional state, we investigated the contribution of NPY signalling to olfactory perception in an odorant preference assay using a food cue associated mixture of amino acids. While the starved fish showed a strong preference for the baited side, recently fed fish were agnostic to the amino acid cue suggesting that the positive valence of food-related odorants is tuned to the prevailing nutritional state. Administration of NPY Y1 receptor antagonist to starved animals also reduced the attraction towards amino acids. Since NPY is a strong orexic agent, this behaviour could also be attributed to the loss of feeding drive following NPY inhibition. We tested this possibility using an amino acid based aversion assay and found that NPY activity was necessary for amino acid perception. As central administration of glucose reduced NPY expression in OE, it should mimic the fed state in its response to amino acids. While glucose injected fish showed no preference for amino acids, exogenous administration of NPY was sufficient to restore the response. Collectively, these experiments implicate a central role for NPY as a mediator of nutritional state-dependent tuning of olfactory perception of food-related cues. To investigate if the differences in olfactory response observed in fed and starved states extend to non-food -associated odorants, we evaluated the response to the alarm substance Schreckstoff (a non-amino acid olfactory cue). Irrespective of their nutritional states, the fish responded similarly to Schreckstoff suggesting that the energy state-dependent modulation is specific to food cues. Teleost fish employ a vomeronasal-like amino acid sensing mechanism, largely confined to microvillar OSNs (Sato *et al.*, 2005; Koide *et al.*, 2009), while Schreckstoff is processed by crypt OSNs (Mathuru *et al.*, 2012). NPY-based neuromodulation appears to be limited to microvillar OSNs. This claim is substantiated by the observation that almost all the amino acid activated OSNs in starved animals (heightened NPY signalling) co-expressed TrpC2 channels. This cell type -dependent partitioning limits NPY-mediated tuning of olfactory sensitivity to amino acids, the major food-associated odorant cue.

Interestingly, we detected little NPY expression in microvillar OSNs. This suggests that NPY, released in a non-synaptic manner from other OSN subtypes and the terminal nerve, acts on NPY receptors on microvillar OSNs and modulates their activity. It remains unclear what triggers the change in NPY expression in response to nutritional states. Although the OB is intrinsically glucosensitive (Fadool *et al.*, 2011; Tucker *et al.*, 2013), we do not know which glucoresponsive source influences the NPY expression.

Modulators like orexin A, insulin and endocannabinoids influence olfactory sensitivity by acting at the level of the OB (Fadool *et al.*, 2000; Apelbaum *et al.*, 2005; Blakemore *et al.*, 2006; Soria-Gomez *et al.*, 2014). The *Drosophila* NPY orthologue, sNPF, has been reported to mediate pre-synaptic facilitation at the first order synapse (Root *et al.*, 2011). However, our evaluation of OSN activation revealed an additional mechanism involving direct modulation of OSN sensitivity to odorants. Amino acid stimulation evoked a strong activation of OSNs in starved animals, but this response was blunted in fed fish. Inhibition of NPY signalling by Y1R antagonist or suppression of endogenous NPY expression by glucose administration resulted in attenuation of OSN activation by amino acids. Conversely, application of exogenous NPY peptide to glucose-injected animals was sufficient to restore the OSN activity. Thus energy-deficient animals are more responsive to odorants. Indeed, OSN activation by amino acids showed a sharp increase in starved animals compared to fed fish. In sum, we show that increased NPY activity lowers the threshold for amino acid detection. Strikingly, this study also revealed a robust co-operative effect between NPY and amino acid signalling that resulted in a switch-like increase in OSN sensitivity to amino acids.

Elevated sNPF signalling in OSNs of starved flies, via increased expression of the sNPF receptor, has been shown to mediate presynaptic facilitation by increasing glutamate release at the OSN – Projection neuron (PN) synapse (Root *et al.*, 2011). Similar observations are reported from *in vitro* studies on olfactory bulb interneurons (Blakemore *et al.*, 2006). While not excluding a mechanism involving pre-synaptic facilitation of neurotransmitter release, our study uncovers an additional mechanism of dynamic adjustment of OSN activation thresholds by NPY signalling. NPY signalling in the peripheral olfactory circuitry is found across invertebrate and vertebrate olfactory lineages (Hansel *et al.*, 2001; Mathieu *et al.*, 2002; Gaikwad *et al.*, 2005; Mousley *et al.*, 2006; Root *et al.*, 2011). Indeed, externally applied NPY increased the electro-olfactogram (EOG) response to amino acids in OE isolated from starved but not fed salamanders and rats (Mousley *et al.*, 2006; Negroni *et al.*, 2012).

However, these studies do not report intrinsic differences in EOG responses between starved and fed animals (Mousley *et al.*, 2006; Negroni *et al.*, 2012). This is possibly because these isolated OE preparations are devoid of endogenous NPY, while an intact behaving animal can access the secreted neuropeptide. The rat study also demonstrated an elevation in NPY receptor expression upon starvation (Negroni *et al.*, 2012). Similar increase in sNPF receptor expression has been reported in starved flies (Root *et al.*, 2011). NPY receptor expression may also increase in the OSNs of starved zebrafish. This increase will augment NPY signalling in these animals along the lines indicated in this study. A recent study in *Drosophila* has reported the modulation of OSN responses by NPF (fly orthologue of NPY) signalling, suggesting the conservation of this modality across phyla (Lee *et al.*, 2017).

Amino acids are typically perceived by microvillar OSNs expressing the vomeronasal class II receptors (V2Rs) that are coupled to PLC activity. PLC activation, in turn, leads to OSN depolarization via the opening of DAG-gated TrpC2 channels (Lucas *et al.*, 2003; Sansone *et al.*, 2014). Zebrafish and *Xenopus*, lack a distinct vomeronasal organ, instead, vomeronasal receptor (VR) expressing OSNs are found in the OE (Sato *et al.*, 2005; Koide *et al.*, 2009; Sansone *et al.*, 2014). We show that OSN activation by amino acid stimulation in zebrafish is also dependent PLC signalling and TrpC2 activity. Importantly, NPY-dependent enhancement of OSN sensitivity is also dependent on PLC activity. Canonical stimulation of NPY receptors involves G_αi_ signalling resulting in inhibition of adenylyl cyclase activity (Herzog *et al.*, 1992). However, heterotrimeric G-proteins can potentially generate bifurcating signals, one mediated by G_αi_ and the other by the G_βγ_ subunits. The latter subunit is known to activate PLC and there is evidence that NPY signalling, in certain contexts, leads to PLC activation (Selbie *et al.*, 1995; Selbie *et al.*, 1997). Thus, NPY and amino acid signalling converge on PLC activation (Figure 6E). This convergence offers a potential mechanism where the additive effect of the two convergent pathways lowers the activation threshold of OSNs (Figure 6E, F). Interestingly, the centrality of PLC activity also offers a potential mechanistic explanation for the co-operative, switch-like increase in microvillar OSN activation observed for co-incident NPY and amino acid signalling. Synergistic activation of PLC isoforms has been extensively reported in studies investigating GPCR signalling cross talk (Zhu & Birnbaumer, 1996; Werry *et al.*, 2003). PLC isoforms, like PLCβ1-3, have two distinct binding sites for G_αq_ and G_βγ_, and simultaneous occupation of these sites can result in synergistic activation (Zhu & Birnbaumer, 1996; Yoon *et al.*, 1999; Philip *et al.*, 2010). In the case of PLCβ3, this cooperative activation has been modeled as a two-state coincidence detector (Philip *et al.*, 2010). NPY signalling, via G_βγ_ activation, has been demonstrated to potentiate adrenergic and purinergic signalling (Selbie *et al.*, 1997). If a similar mechanism were operational in zebrafish OSNs, then it would be expected that the amino acid signalling via V2Rs would involve G_αq_ activity. We tested this possibility and found that GP2A treatment (a specific G_αq_ activity blocker) prevented amino acid mediated OSN activation. This observation offers new insight into olfactory processing of amino acid cues in fish. It is worth noting that while G_αi_ and G_αo_ are predominantly implicated in mammalian VNO transduction, there is also available evidence suggesting the involvement of G_αq_ (Wekesa *et al.*, 2003; Thompson *et al.*, 2006).

These experiments offer a mechanistic framework for NPY-mediated lowering of activation threshold in amino acid sensing OSNs, where the level of NPY signalling tunes the baseline PLC activity in the neurons (Figure 6F). Low NPY signalling (in recently fed or glucose treated fish) results in low basal levels of PLC and the odorant triggered PLC activation has to be substantial to activate the neuron. Elevated NPY signalling in starved animals, along with coincident signalling from amino acid stimulated V2Rs, converges onto PLC. This convergence augments PLC activity, possibly cooperatively, resulting in switch-like lowering of the threshold necessary to stimulate TrpC2 channels in order to activate the OSN.

Our study offers ethologically relevant evidence that in vertebrates NPY-signalling is responsive to internal energy states and tunes olfactory sensitivity to food-related odorants. We uncover a novel mechanism involving NPY signalling dependent peripheral modulation of OSN activation thresholds via PLC activity as a central component of tuning olfactory responses to nutritional states.

## ETHICS APPROVAL AND CONSENT TO PARTICIPATE

All protocols used in this study were approved by Institutional Animal Ethics Committee.

## CONSENT FOR PUBLICATION

Not applicable.

## AVAILABILITY OF DATA AND MATERIAL

All data generated or analysed during this study are included in this published article. The raw data are available from the corresponding author on reasonable request.

## AUTHOR CONTRIBUTIONS

Conceptualization, N.S. and A.G.; Investigation, T.K., A.D., A.M., Ak.G., Ar.M; Writing, N.S. and A.G.; Funding Acquisition, A.G.

## ACKNOWLEDGMENTS

Science and Engineering Research Board (SERB), Govt. of India grant [SB/SO/AS-038/2013] to AG and intramural funding from IISER Pune, supported this work. The authors acknowledge the IISER Pune Microscopy Facility and the National Facility for Gene Function in Health and Disease (NFGFHD) at IISER Pune for access to equipment and infrastructure.

